# Arsenic is a potent co-mutagen of ultraviolet light

**DOI:** 10.1101/2023.02.22.529578

**Authors:** Rachel M. Speer, Shuvro P. Nandi, Karen L. Cooper, Xixi Zhou, Hui Yu, Yan Guo, Laurie G. Hudson, Ludmil B. Alexandrov, Ke Jian Liu

**Affiliations:** Department of Pharmaceutical Sciences, College of Pharmacy, University of New Mexico, Albuquerque, NM 87106, USA; Department of Cellular and Molecular Medicine, UC San Diego, La Jolla, CA, 92093, USA; Moores Cancer Center, UC San Diego, La Jolla, CA, 92037, USA; Department of Bioengineering, UC San Diego, La Jolla, CA, 92093, USA; Department of Internal Medicine, Division of Molecular Medicine, University of New Mexico, Albuquerque, NM 87106, USA; Stony Brook Cancer Center, Stony Brook University, Stony Brook NY 11794, USA; Department of Pathology, Stony Brook University School of Medicine, Stony Brook, NY 11794, USA

**Author notes:** These authors contributed equally to this work (Rachel M. Speer, Shuvro P. Nandi).

## Abstract

Environmental co-exposures are widespread and are major contributors to carcinogenic mechanisms. Two well-established environmental agents causing skin cancer are ultraviolet radiation (UVR) and arsenic. Arsenic is a known co-carcinogen that enhances UVR’s carcinogenicity. However, the mechanisms of arsenic co-carcinogenesis are not well understood. In this study, we utilized primary human keratinocytes and a hairless mouse model to investigate the carcinogenic and mutagenic properties of co-exposure to arsenic and UVR. *In vitro* and *in vivo* exposures revealed that, on its own, arsenic is neither mutagenic nor carcinogenic. However, in combination with UVR, arsenic exposure has a synergistic effect leading to an accelerated mouse skin carcinogenesis as well as to more than 2-fold enrichment of UVR mutational burden. Notably, mutational signature ID13, previously found only in UVR-associated human skin cancers, was observed exclusively in mouse skin tumors and cell lines jointly exposed to arsenic and UVR. This signature was not observed in any model system exposed purely to arsenic or purely to UVR, making ID13 the first co-exposure signature to be reported using controlled experimental conditions. Analysis of existing genomics data from basal cell carcinomas and melanomas revealed that only a subset of human skin cancers harbor ID13 and, consistent with our experimental observations, these cancers exhibited an elevated UVR mutagenesis. Our results provide the first report of a unique mutational signature caused by a co-exposure to two environmental carcinogens and the first comprehensive evidence that arsenic is a potent co-mutagen and co-carcinogen of UVR. Importantly, our findings suggest that a large proportion of human skin cancers are not formed purely due to UVR exposure but rather due to a co-exposure of UVR and other co-mutagens such as arsenic.

## INTRODUCTION

Carcinogens are agents that result in cancer formation^1^, with many carcinogens causing cancer by directly generating somatic mutations^2^. Recent experimental studies have also unambiguously described non-mutagenic carcinogens, where cancers were induced in mice by exposing them to suspected human carcinogens without observing an elevation in somatic mutations^3^. Further, prior studies have provided evidence for the existence of co-carcinogens, which are agents that are not carcinogenic on their own, but they rather promote the effects of other carcinogens^4^. Lastly, limited prior evidence has been offered for co-mutagenic agents, which are generally non-mutagenic but, in combination with another agent, can have synergistic effect leading to a highly accelerated mutagenesis^5^.

Arsenic is a naturally occurring element and a known environmental contaminant found in high concentrations in drinking water within the United States and across the world, particularly from water sourced from wells^6,7^. The International Agency for Research on Cancer has classified arsenic as carcinogenic to humans based on strong evidence linking arsenic exposure to cancers of the lung, bladder, kidney, and skin^8,9^. While skin cancer is commonly associated with exposure to ultraviolet radiation (UVR) from sunlight, arsenic is a known co-carcinogen of UVR^10-12^. Further, epidemiological studies have shown an increased cancer risk for developing UVR-associated skin cancer in populations exposed to high-levels of arsenic and these results have been supported by experimental studies^13-16^. While prior research has shown arsenic inhibits repair of UVR-induced DNA damage^17-19^ the mutagenic properties of arsenic co-exposure have not been well understood.

Analysis of mutational signatures allows elucidating the mutagenic processes that lead to cancer^20^. Previously, we and others have described more than 100 different mutational signatures including ones associated with environmental carcinogens, failure of DNA-repair pathways, infidelity of replicating polymerases, chemotherapeutics, and many others^21,22^. Only one study has investigated the mutational patterns of arsenic in human cancer by inconclusively examining a single never-smoker lung cancer patient chronically exposed to arsenic^23^. Further, while induced pluripotent stem cell lines have been exposed to arsenic, no arsenic mutational signature was found^24^. In contrast, exposure to UVR from sunlight is known to induce specific DNA damage and several distinct UVR-associated mutational signatures have been identified in human tumors, normal human tissues, and experimental systems^25-29^.

Notably, mutational signatures of single base substitutions (SBSs), termed COSMIC signatures SBS7a/b/c/d, have been found at extremely high levels in most cancers of the skin^22^ as well as in experimental systems exposed to UVR^24^. SBS7a/b are characterized by C>T mutations at dipyrimidines and have been associated with DNA damage due to UVR, including both 6,4-photoproducts and cyclobutane pyrimidine dimers (CPDs)^30-32^. Signatures SBS7c and SBS7d are characterized by T>A and T>C mutations, respectively, and while these UVR signatures are exclusively found in cancers of the skin, their etiology remains mysterious^22,32,33^.

A doublet-base substitution (DBS) signature, termed DBS1, has also been found at high levels in human skin tumors, normal human skin tissues, and experimental systems exposed to UVR^25-29^. Signature DBS1 exhibits almost exclusively CC>TT mutations and it has been attributed to mis-replication of CPDs^22^. Additionally, a mutational signature of small insertions and deletions (indels), termed, COSMIC signature ID13, has been found exclusively in cancers of the skin in sun exposed areas and, thus, it has been attributed to exposure to ultraviolet light^22,33^. ID13 exhibits a particular pattern that includes a deletion of a single thymine at a thymine-thymine dimer^33^.

In this study, we leverage well controlled *in vitro* and *in vivo* co-exposures in combination with whole-genome sequencing and mutational signatures analysis to investigate the carcinogenic and mutagenic properties of arsenic and solar-simulated UVR co-exposure. Our experimental findings reveal that, in combination with UVR, arsenic exposure has a synergistic effect leading to an enhanced mouse skin carcinogenesis as well as to more than 2-fold enrichment of UVR mutational burden. Importantly, signature ID13 is uniquely due to arsenic and UVR co-exposure and, comparisons with genomics data from previously generated skin cancers demonstrate that ID13 is found exclusively in a large proportion of human skin cancers with an elevated UVR mutagenesis. Our results demonstrate that arsenic is a potent co-mutagen of ultraviolet light that amplifies UVR mutagenesis and that generates a unique mutational signature commonly found in human cancers of the skin.

## RESULTS

### In vitro and in vivo experimental designs

To examine the mutagenic properties of co-exposures to arsenic (As) and UVR, we used both *in vitro* and *in vivo* models. Specifically, an immortalized keratinocyte cell line, N/TERT1^34^, was utilized under the following conditions: *(i)* no treatment (NT); *(ii)* irradiation with UVR (3 kJ/m^2^); *(iii)* treatment with arsenic (1 μM); and *(iv)* pre-treatment with arsenic (1 μM) for 24 hours followed by irradiation with UVR (3 kJ/m^2^). Arsenic exposures were continued for 24 hours post-UVR irradiation during the time in which UVR generated DNA damage is likely being repaired. All cells were cultured for additional 24 hours after their respective exposures and, subsequently, subjected to barrier bypass-clonal expansions^35^ and whole-genome sequencing (**Fig. 1*a***). The selected arsenic and UVR exposure levels align with previous studies investigating arsenic-UVR co-carcinogenesis^36-38^. In most cases, the combined exposure of arsenic and UVR resulted in similar levels of cytotoxicity to the ones due to UVR exposure alone (**Fig. 1*b***); cytotoxicity was measured relative to the NT group. Consistent with conditions used in previous evaluation of environmental carcinogens^24^, we clonally expanded cells from exposures resulting in approximately 50% relative cell death (**Fig. 1*b***).

**Figure 1.**
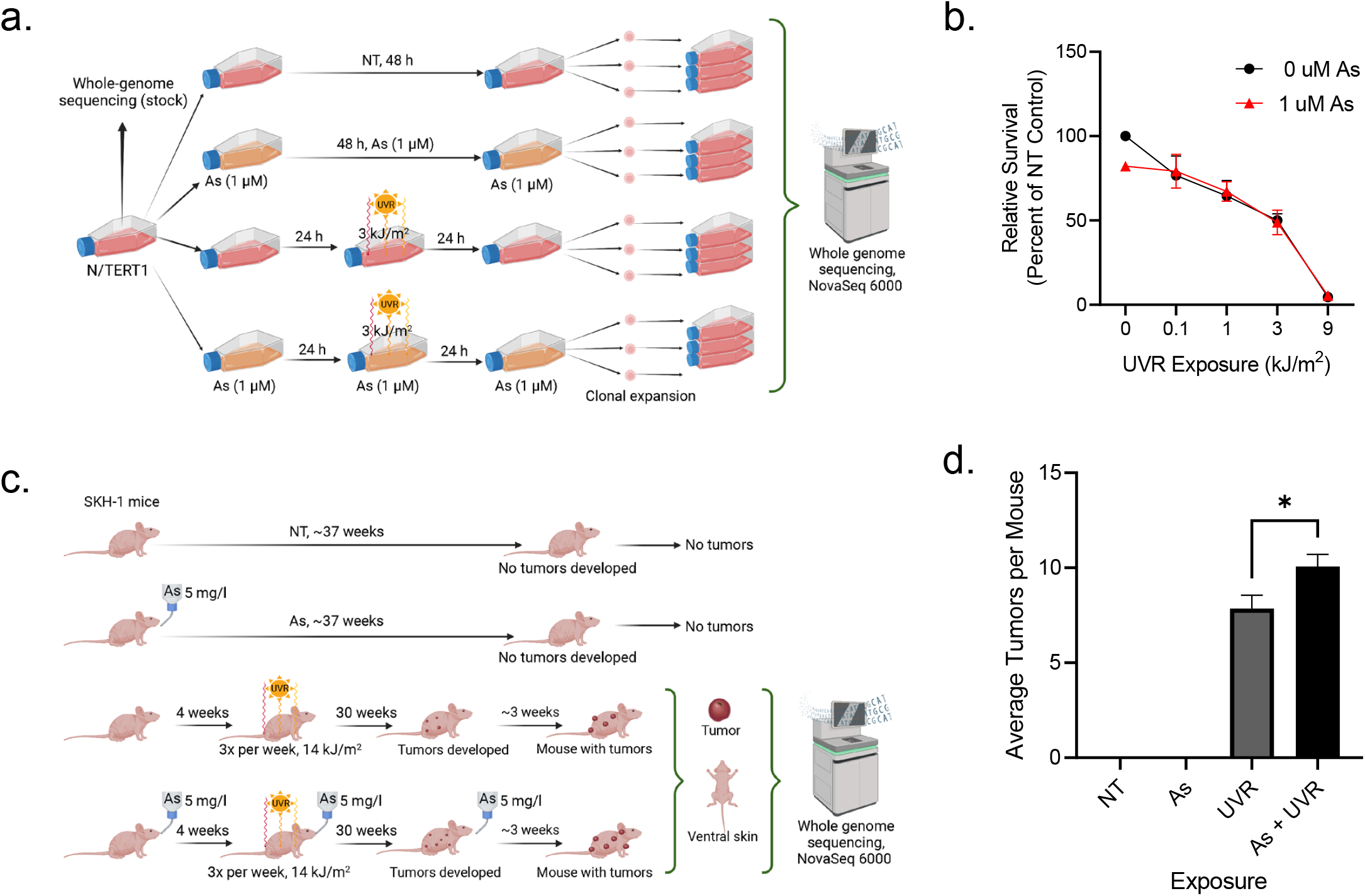
Experimental design for mutational signatures co-exposure analysis. **a**. Experimental design for N/TERT1 cells, where somatic mutations were called from clones expanded from treatment groups, no treatment (NT) control, arsenic (As), ultraviolet light radiation (UVR), and As plus UVR, against the bulk sequenced stock. **b**. Y-axis reflects the relative survival of exposed cells measured using the percentage of clonogenic survival compared to survival of the NT control. The x-axis corresponds to the total amount of energy delivered per unit area. The red line depicts survival of cells pre-treated with arsenic (1 μM), while the black line reflects survival of cells without arsenic pre-treatment. No statistically significant difference in survival were observed in cells pre-treated with arsenic and cells without arsenic pre-treatment (two-sided t-test; *n*=2 derived from 2 independent experiments with technical replicates each for all experimental conditions). The experimental conditions used in this study utilized exposure levels leading to 50% relative survival in N/TERT1 cells in alignment with previously published studies^24^. **c**. Experimental design utilizing SKH-1 hairless mouse model, where mice were separated into four groups, including: a NT group; arsenic exposed group; UVR exposed group; and As plus UVR exposed group. No tumors developed in NT or arsenic alone groups. Somatic mutations in skin tumors from UVR as well as As plus UVR exposed mice were identified by comparing the sequenced tumor tissues to the sequenced ventral (non-UVR exposed) normal skin from the same animal. **d**. Y-axis reflects the average number of tumors per mouse, while the x-axis corresponds to the different experimental conditions. The tumorigenicity for the mouse model shows arsenic significantly enhances tumor burden in UVR exposed mice (p-value<0.05; two-sided t-test; *n*=14 for all conditions). Data represent the mean ± SEM. Statistical details are reported in the **Methods** section.

To confirm the observed *in vitro* results, we also utilized a SKH-1 hairless mouse model^39^ where mice were separated into four groups (*n*=14 for each group), including: *(i)* a NT group; *(ii)* a group where mice were exposed to arsenic in their drinking water (5 mg/l); *(iii)* a group where mice were exposed to UVR three times per week (14 kJ/m^2^); and *(iv)* a group where mice were exposed to arsenic in their drinking water (5 mg/l) and exposed to UVR three times per week (14 kJ/m^2^; **Fig. 1*c***). Mice are faster metabolizers of arsenic compared to humans, and thus, higher arsenic concentrations are required to induce responses similar to those that would be seen in humans at lower concentrations^40,41^. No overt toxicity was observed in mice exposed to 5 mg/l. The UVR exposure chosen is approximately half the minimal exposure level that results in erythema (reddening of the skin) and is therefore relevant to environmental exposures. No tumors developed in mice in the NT or arsenic alone groups, however, tumors developed in the UVR group and tumor burden was 1.3-fold enhanced by co-exposure with arsenic (p-value: 0.0265; two-sided t-test; **Fig. 1*d***).

### Arsenic affects UVR mutagenesis in vitro

Somatic mutations were identified from all whole-genome sequenced N/TERT1 cells by bioinformatically comparing them to the whole-genome sequenced stock cells (**Methods**; **Fig. 1*a***). Statistical comparisons for the N/TERT1’s mutational landscapes were performed amongst controls and the three different types of exposures using one-way ANOVA with Tukey post hoc correction for multiple comparisons (**Fig. 2*a&c***). Arsenic alone did not increase the total numbers of SBSs, DBSs, or indels in N/TERT1 cells when compared to the ones found in NT controls (**Fig. 2*a***). In contrast, UVR exposure resulted in a significant increase of 3.9-fold for SBSs and 10-fold for DBSs when compared to NT controls (p-values: 0.0040 and 0.0345, respectively). C>T, T>C, C>A, T>A, and T>G substitutions were significantly elevated when compared to their levels in untreated controls (p-values: 0.0074, 0.0476, 0.0152, 0.0459, and 0.0309, respectively; **Supplementary Fig. 1*a-d***). Importantly, samples co-exposed to arsenic and UVR exhibited approximately 1.8- and 2.1-fold significant enrichment of SBSs and DBSs, respectively, when compared to samples exposed to UVR alone (p-values<0.05). Specifically, C>T mutations contributed most mutations in UVR exposed cells and arsenic co-exposure resulted in 2.2-fold increase of these mutations compared to UVR alone (p-value: 0.001; **Supplementary Fig. 1*a***). Arsenic also significantly increased C>G mutations compared to UVR alone (p-value: 0.0112).

**Figure 2.**
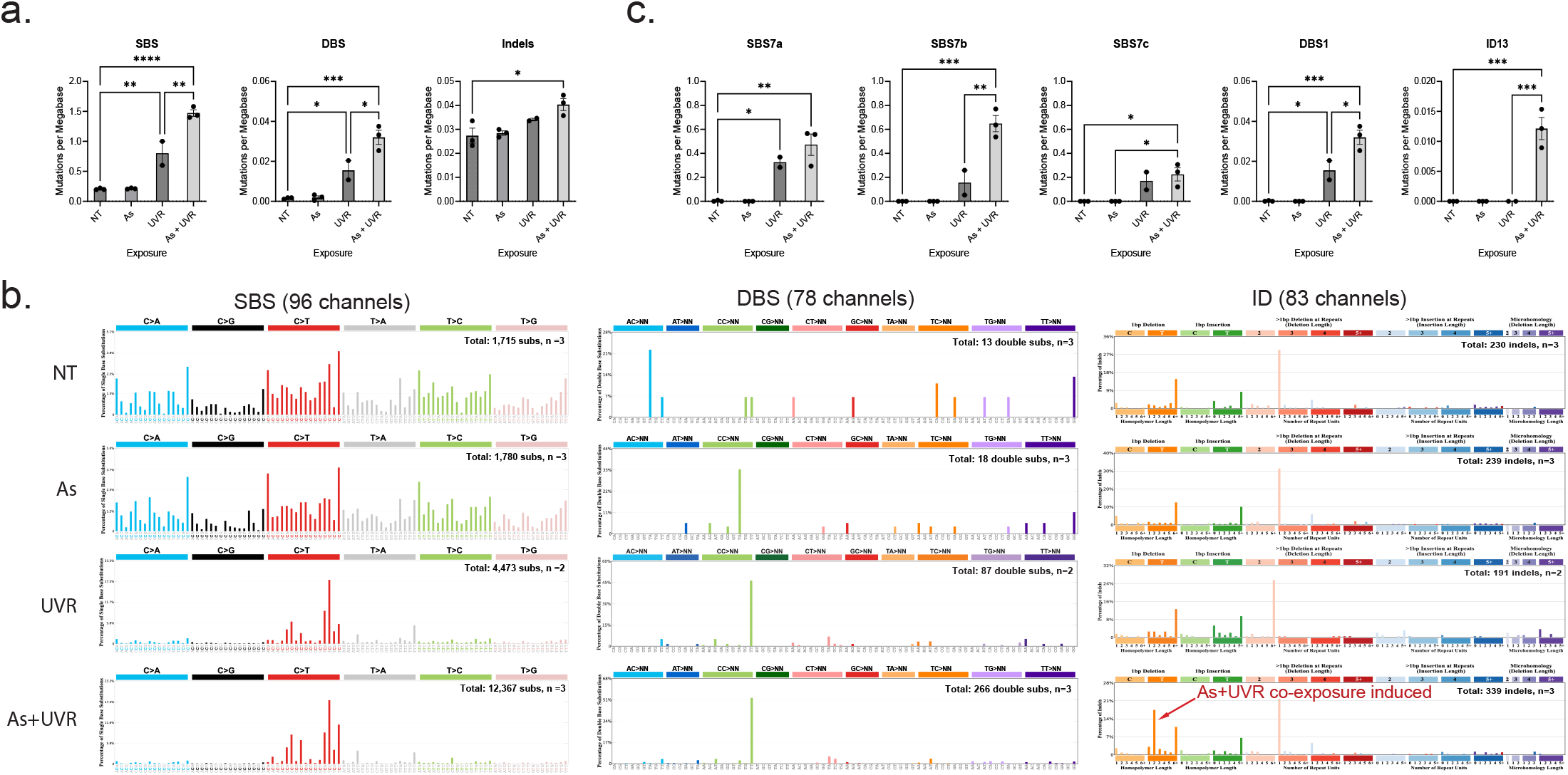
Arsenic enhances somatic mutations imprinted by ultraviolet light in N/TERT1 cells. **a**. Y-axes reflect the amounts of substitutions (SBS; left), doublets (DBS; middle), or small insertions and deletions (Indels; right) measured in somatic mutations per megabase. X-axes correspond to the different types of exposures including: no treatment (NT) control, arsenic (As), ultraviolet light radiation (UVR), and As plus UVR. Bar plots represent the mean ± SEM; individual biological replicates are shown as black circles. **b**. Patterns of single base substitutions (SBS) are shown on the left using the SBS-96 classification scheme^44^ on the x-axes. Patterns of doublet base substitutions (DBS) are shown in the middle using the DBS-78 classification scheme^44^ on the x-axes. Patterns of small insertions and deletions (ID) are shown on the right using the ID-83 classification scheme^44^ on the x-axes. Each plot represents the average mutational profile of each treatment group across all samples in that group. Y-axes are scaled differently in each plot to optimally show each average mutational pattern with the y-axes reflecting the percentage of mutations for the respective mutational scheme. **c**. Y-axes reflect the amounts of COSMIC mutational signatures measured in somatic mutations per megabase. X-axes correspond to the different types of exposures. Bar plots represent the mean ± SEM; individual biological replicates are shown as black circles. Significance was evaluated using one-way ANOVA with Tukey’s multiple comparisons test; *n*=3 for NT, As, and As plus UVR and *n*=2 for UVR alone derived from independent clones. *p-value<0.05, **p-value<0.01, ***p-value<0.005, ****p-value<0.001. Statistical details are reported in the **Methods** section.

The total number of indels was 1.5-fold elevated in samples co-exposed to arsenic and UVR when compared to NT control (p-value: 0.0177) but not compared to UVR alone (**Fig. 2*a***).

The mutational patterns of arsenic exposed cells were identical to the ones of non-treated controls for both substitutions and indels (cosine similarity: 0.97; **Fig. 2*b***). The numbers of doublets were too few to perform a comparison between NT controls and arsenic exposed cells. In contrast, UVR-exposed N/TERT1 cells exhibited a distinct pattern of C>T substitutions at dipyrimidines as well as high levels of CC>TT doublets (**Fig. 2*b***). Further, the mutational patterns of both substitutions and doublets were remarkably similar between cells exposed only to UVR and cells exposed jointly to arsenic and UVR (cosine similarity: 0.97). Nevertheless, the pattern of small insertions and deletions was different between these two exposures with a striking elevation of single thymine deletions at a thymine-thymine dimers in the cells exposed to arsenic plus UVR (**Fig. 2*b***). Additionally, an examination of previously generated datasets^24^ revealed that the substitution patterns of UVR in N/TERT1 cells are similar to the ones observed in human induced pluripotent stem cells (iPSCs) exposed to UVR (cosine similarity: 0.96). In contrast, a distinct difference can be seen in the iPSC indel profile which lacks the thymine deletion at thymine-thymine dimers observed in arsenic and UVR co-exposed N/TERT1 cells (cosine similarity: 0.35). Overall, the indel profile of UVR-exposed iPSC cells was consistent with the one of UVR-exposed N/TERT1 cells and neither UVR-exposed iPSC cells nor UVR-exposed N/TERT1 cells harbored the unique indel pattern observed in N/TERT1 cells co-exposed to arsenic and UVR.

Analysis of COSMIC mutational signatures revealed that three of the four UVR-associated SBS signatures, SBS7a/b/c, as well as the UVR-associated signatures DBS1 and ID13 were found in N/TERT1 cells exposed to UVR (**Fig. 2*c***). No UVR-associated signatures were identified in untreated N/TERT1 cells or in N/TERT1 exposed purely to arsenic. Co-exposure to arsenic resulted in a 4.2-fold increase of SBS7b and 2.1-fold increase of DBS1 (p-value: 0.0015 and 0.0143, respectively) but not to a statistically significant elevation of signatures SBS7a or SBS7c compared to UVR alone. Remarkably, signature ID13 was exclusively identified in the N/TERT1 cells co-exposed to arsenic and UVR but not in cells exposed purely to UVR (p-value: 0.0005; **Fig. 2*c***). Consistent with the analysis of COSMIC mutational signatures, analysis of *de novo* signatures revealed an elevation of SBS signatures as well as an indel signature, resembling ID13, found exclusively in samples co-exposed to UVR and arsenic (**Supplementary Fig. 2**).

### Arsenic affects UVR mutagenesis in vivo

The performed *in vitro* exposures were complemented by almost identical exposures in a SKH-1 hairless mouse model (**Fig. 1*c***). No tumors were observed in the NT group (*n*=14) or in mice exposed to arsenic alone (*n*=14; **Fig. 1*d***). For a subset of UVR-exposed mice, a tumor and matched normal skin tissue from the ventral (non-UVR exposed) side of each animal were whole-genome sequenced and, subsequently, bioinformatically compared to derive somatic mutations in the tumor tissue (**Methods**; **Fig. 1*c***). Statistical comparisons between the mutational landscapes of the tumors in mice exposed to UVR alone and the tumors in mice co-exposed to arsenic and UVR were performed using FDR-corrected two-sided t-tests (**Methods**).

Tumors from mice co-exposed to arsenic and UVR exhibited approximately 6-fold enrichment of substitutions, 6-fold enrichment of doublets, and 3-fold enrichment of indels when compared to tumors from mice only exposed to UVR (q-values: 0.0009, 0.0009, 0.0009, respectively; **Fig. 3*a***). Similar to N/TERT1 cells, all types of single base substitutions and doublet base substitutions were significantly elevated in tumors due to co-exposure to arsenic and UVR (q-values<0.05; **Supplementary Fig. 1*e-h***). Further, both insertions and deletions were found to be increased approximately 3-fold in tumors from co-exposed mice when compared to tumors due to UVR alone (q-values: 0.0014 and 0.0023, respectively; **Supplementary Fig. 1*g-h***).

**Figure 3.**
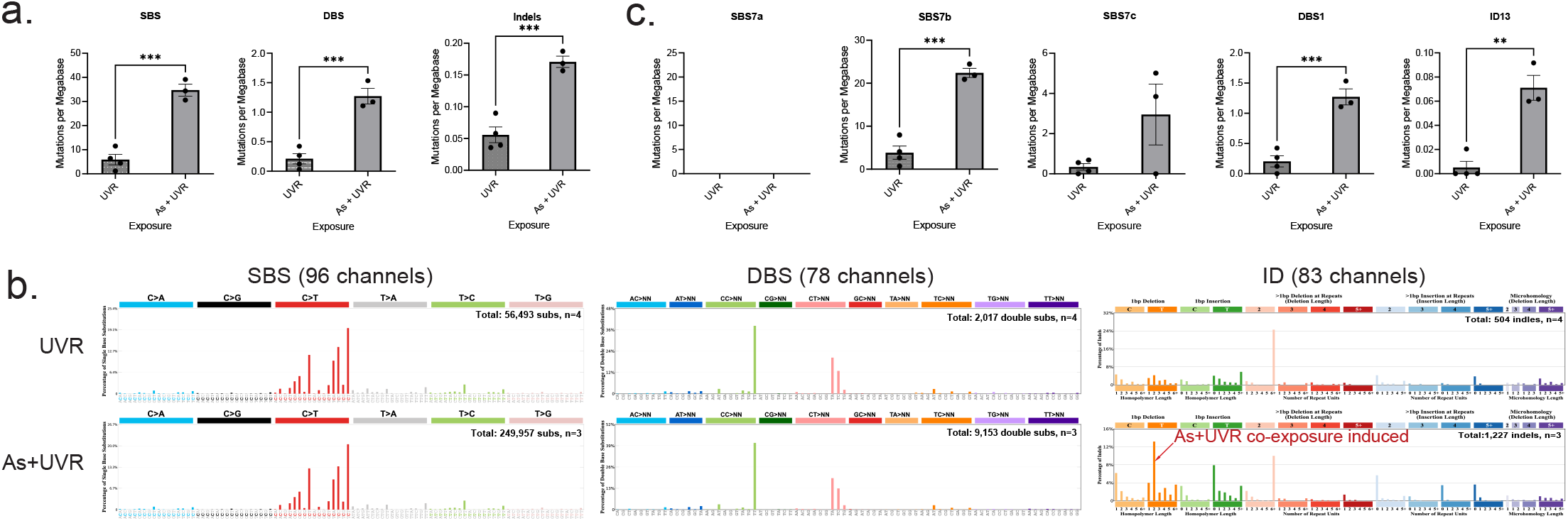
Arsenic enhances somatic mutations imprinted by ultraviolet light in skin cancers from SKH-1 hairless mice. **a**. Y-axes reflect the amounts of substitutions (SBS; left), doublets (DBS; middle), or small insertions and deletions (Indels; right) measured in somatic mutations per megabase. X-axes correspond to the different types of exposures including: ultraviolet light radiation (UVR), and arsenic (As) plus UVR. Bar plots represent the mean ± SEM; individual biological replicates are shown as black circles. **b**. Patterns of single base substitutions (SBS) are shown on the left using the SBS-96 classification scheme^44^ on the x-axes. Patterns of doublet base substitutions (DBS) are shown in the middle using the DBS-78 classification scheme^44^ on the x-axes. Patterns of small insertions and deletions (ID) are shown on the right using the ID-83 classification scheme^44^ on the x-axes. Each plot represents the average mutational profile of each treatment group across all samples in that group. Y-axes are scaled differently in each plot to optimally show each average mutational pattern with the y-axes reflecting the percentage of mutations for the respective mutational scheme. **c**. Y-axes reflect the amounts of COSMIC mutational signatures measured in somatic mutations per megabase. X-axes correspond to the different types of exposures. Bar plots represent the mean ± SEM; individual biological replicates are shown as black circles. Significance was evaluated using FDR corrected two-sided t-tests; *n*=4 for UVR alone and *n*=3 for As plus UVR derived from individual animals. **q-value<0.01, ***q-value<0.005. Bar plots represent the mean ± SEM; individual replicate values are shown as black circles. Statistical details are reported in the **Methods** section.

A distinct pattern of C>T substitutions at dipyrimidines was observed in all mouse tumors (**Fig. 3*b***). The pattern was identical between tumors due to UVR alone and tumors due to arsenic and UVR (cosine similarity: 0.99; **Fig. 3*b***). Interestingly, this pattern of single base substitutions is also similar to previous data from mouse cell lines^27^ exposed to UVR (cosine similarity: 0.98) while differing from the substitution patterns observed in N/TERT1 cells (cosine similarity: 0.84) or in UVR-associated human skin cancers (cosine similarity: 0.80)^25^. Specifically, UVR-imprinted patterns in mouse tumors and mouse cell lines have a distinctly high peak of C>T mutations at Tp<u>T</u>pT trinucleotides (mutated base underlined; **Fig. 3*b***) which is absent in human tumors^27^ or in human cell lines (**Fig. 2*b***). Similarly, CC>TT and CT>NN dinucleotides were observed in all UVR-associated mouse tumors (**Fig. 3*b***). The CC>TT mutational pattern in mouse tumors was similar to the one observed in N/TERT1 cells (**Fig. 2*b***) but the CT>NN dinucleotides were unique for mouse tumors and mouse cell lines^27^ and CT>NN dinucleotides have not been found in human skin cancers or in UVR-exposed human cell lines. Importantly, similar to human cell lines, the mouse tumors exhibited a striking elevation of single thymine deletions at thymine-thymine dimers in the cells co-exposed to UVR and arsenic (**Fig. 3*b***).

Evaluating the COSMIC mutational signature in the UVR-exposed mouse tumors elucidated the presence of signatures SBS7b, SBS7c, DBS1, and ID13. Consistent with the *in vitro* observations, signatures SBS7b and DBS1 were almost 6-fold enriched in the tumors due to co-exposure to arsenic and UVR (q-values: 0.0006 and 0.001, respectively; **Fig. 3*c***). Further, as in the cell line experiments, signature ID13 was exclusively identified in tumors due to co-exposure to UVR and arsenic but not in tumors purely due to UVR exposure (**Fig. 3*c***). Consistent with the analysis of COSMIC mutational signatures and the observations in N/TERT1 cell lines, analysis of *de novo* signatures from mouse tumors revealed an elevation of SBS signatures and an indel signature, resembling ID13, highly elevated in samples co-exposed to UVR and arsenic (**Supplementary Fig. 2**).

### Evaluating arsenic-like co-exposures in human skin cancer

Our experimental results revealed that ID13 is generated exclusively in samples jointly co-exposed to arsenic and UVR (**Figs. 2** and **3**). To the best of our knowledge, this study is the first to report ID13 in any experimental system likely due to prior studies focusing purely on UVR exposure without any additional co-exposures^27^. Importantly, ID13 was not observed in any sample exposed purely to UVR (**Figs. 2** and **3**), indicating that a co-exposure to arsenic or to another co-mutagen with similar properties is required for generating signature ID13.

Next, we interrogated 205 previously published^42^ whole-exome sequenced basal cell carcinomas (BCCs) and utilized signature ID13 as a biomarker of potential co-exposure to UVR and arsenic (or another arsenic-like agent). Mutational signature analysis revealed that 19% of BCC samples exhibited ID13 while no evidence for ID13 was found in the remaining samples. The SBS, DBS, and indel mutational profiles of BCCs were partitioned into ID13 negative and ID13 positive samples (**Fig. 4*a***). The SBS and DBS patterns were identical between ID13 negative and ID13 positive BCC sample (cosine similarities >0.98; **Fig. 4*a***). Furthermore, consistent with our experimental results that samples co-exposed to arsenic and UVR exhibited a much higher burden of single and doublet substitutions, the BCC samples harboring ID13 exhibited 1.33-fold elevation of both single base substitutions and doublet base substitutions (q-values<0.05; **Fig. 4*b***). Identical analysis of mutational signature applied to 107 whole-genome sequenced melanomas^43^ from the Pan-Cancer Analysis of Whole Genomes (PCAWG) study yielded similar results with 58% of melanoma genomes harboring ID13 and exhibiting a highly elevated mutational burden of single and doublet substitutions (**Supplementary Fig. 4**).

**Figure 4.**
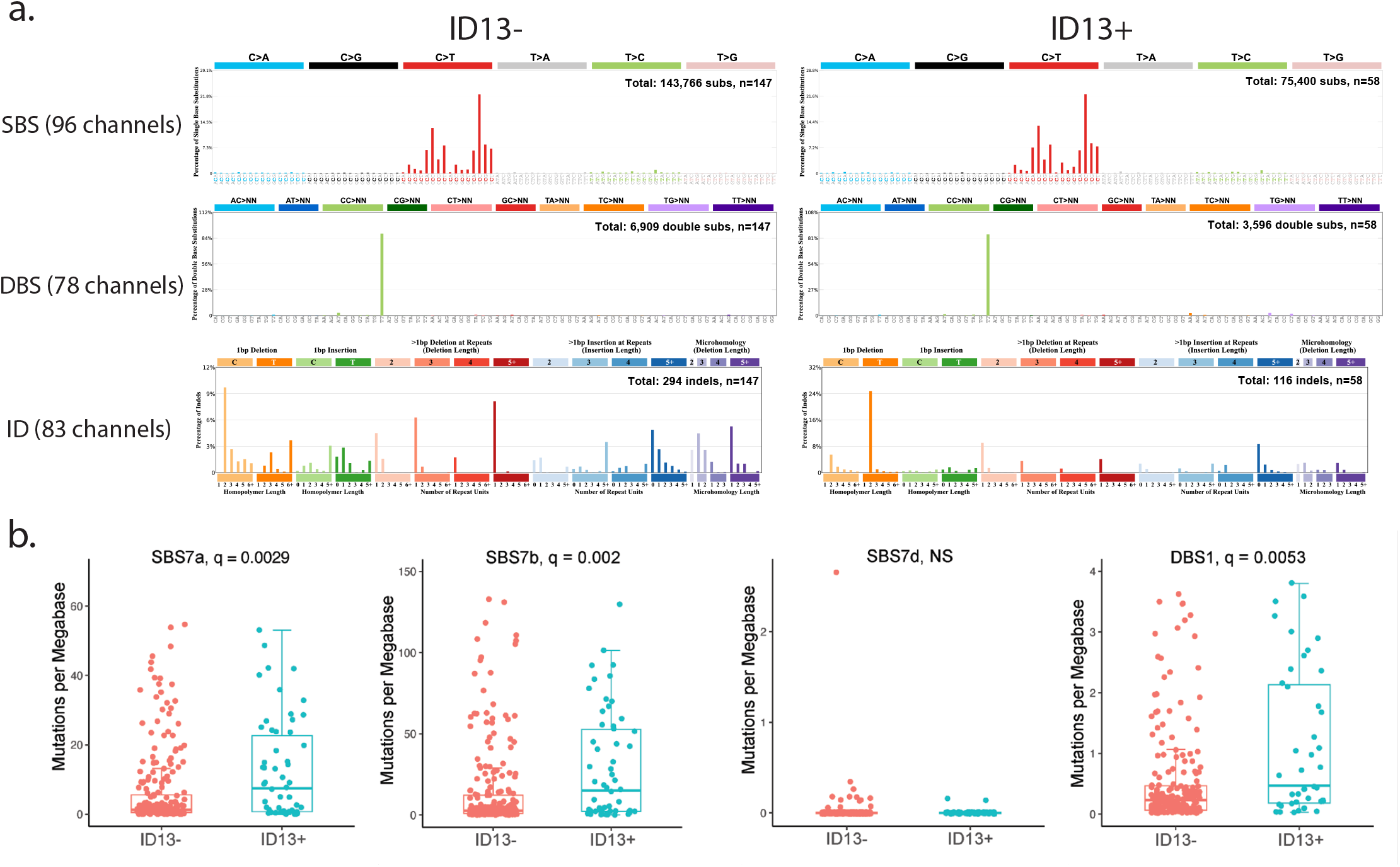
An evaluation of UVR and arsenic-like co-exposure in human basal cell carcinomas. **a**. Patterns of single base substitutions (SBS) in basal cell carcinomas (BCCs) are shown on the left using the SBS-96 classification scheme^44^ on the x-axes. Patterns of doublet base substitutions (DBS) in BCCs are shown in the middle using the DBS-78 classification scheme^44^ on the x-axes. Patterns of small insertions and deletions (ID) in BCCs are shown on the right using the ID-83 classification scheme^44^ on the x-axes. Each plot represents the average mutational profile of each treatment group across all samples in that group. Y-axes are scaled differently in each plot to optimally show each average mutational pattern with the y-axes reflecting the percentage of mutations for the respective mutational scheme. **b**. Mutations per megabase attributed to COSMIC mutational signatures operative in basal cell carcinomas. Each dot reflects the mutations per megabase attributed to each COSMIC signature in each sample. The bounds of the boxplots represent the interquartile range divided by the median, and Tukey-style whiskers extend to a maximum of 1.5 × interquartile range beyond the box. Statistically significant results from FDR corrected two-sided t-tests tests are denoted as q-values. In both panels, basal cell carcinomas were separated on samples containing ID13 (*n* = 58) and basal cell carcinomas without any ID13 (*n* = 147).

## DISCUSSION

In this study, we applied mutational signatures analysis to whole-genome sequencing data from well-controlled *in vitro* and *in vivo* experimental systems to elucidate the carcinogenic and mutagenic potentials of arsenic and ultraviolet light. As expected, the mutational patterns found on the genomes of UVR-exposed cell lines and on mouse cancers were consistent with the set of known UVR mutational signatures derived from human skin cancers. Exposing cell lines purely to environmental relevant concentration of arsenic neither caused an elevated mutational burden nor a specific mutational signature. Further, mice exposed only to arsenic did not develop any tumors. Nevertheless, the co-exposure to arsenic and UVR resulted in an enhanced carcinogenesis and a synergistic elevation of UVR mutagenesis. Importantly, a specific mutational signature, ID13, was found exclusively in samples co-exposed to arsenic and UVR. To the best of our knowledge, this is the first report of a unique mutational signature caused by a co-exposure to two environmental carcinogens and the first comprehensive evidence that arsenic is a potent co-mutagen of UVR.

Our examination of human skin cancers revealed that 19% of basal cell carcinomas and 58% of human melanomas harbor signature ID13 in their genomes. This is a striking result as, based on our experimental data and previous experimental interrogations^24,27^, exposure to UVR alone cannot induce signature ID13. Further, consistent with these experimental findings, human skin cancers with ID13 exhibited an elevated burden of single and doublet substitutions, thus, further implicating an additional co-exposure. Overall, our results suggest that a large proportion of human skin cancers are formed due to co-exposure of UVR and arsenic or due to co-exposure of UVR and another co-carcinogenic agent with similar co-mutagenic properties to the ones of arsenic.

## METHODS

### Cell culture

An hTERT immortalized non-cancerous human keratinocyte cell line (N/TERT1) was used in this study. N/TERT1 cells were established from a male neonate^34^ and respond similarly as primary keratinocytes under experimental conditions^45^. N/TERT1 cells were cultured as monolayers with serum-free DermaLife K Keratinocyte Medium (Lifeline Cell Tech) supplemented with DermaLife K LifeFactors in a humidified incubator at 37°C and 5% CO_2_. Cells were sub-cultured with 0.05/0.02% Trypsin/EDTA (Lifeline Cell Tech) every 3-5 days. Although N/TERT1 is a clonally derived cell line, these cells were established >20 years ago. Through normal passaging, single nucleotide variants arise creating a heterogenous population. Therefore, to reduce noise in the mutational signatures data N/TERT1 were subcloned and a single clone (the grandparent clone) was selected for seeding all experiments. Cells were routinely tested to be negative for mycoplasma and screened for chromosome stability.

### Ultraviolet Radiation (UVR) Exposure

Solar simulated UVR (UVR) exposures were performed using an Oriel 1600 Watt Solar Ultraviolet Simulator (Oriel Corp., Stratford, CT). This solar simulator produces a high intensity UVR beam in both the UVA (320-400 nm) and UVB (280-320 nm) spectrum with an emission ratio of 13:1 (UVA:UVB). The proportion and intensity of UVA/UVB was measured using an ILT2400 radiometer equipped with UVA (SED033), UVB (SED240) and UVC (SED270) detectors (International Light Technologies; Peabody, MA). In vivo exposures were at 14 kJ/m^2^ providing approximately 0.5 minimum erythema dose (MED). Measurements were made with Erythema UV and UVA intensity meter (Solar Light Co., Inc., Philadelphia, PA) to estimate MED. Animal UVR dosing was conducted in groups of 4 - 6 with animals allowed to move freely within the exposure enclosure. Cells and animals were kept in the dark during transport to and from the UVR exposure lamp.

### Cell exposures

Arsenic stock solutions of inorganic arsenic as sodium meta-arsenite (purity >99%; Fluka Chemie) were prepared in double distilled water and filtered through a 0.2 μM filter. For all experiments cells were seeded and allowed to rest for 48 hours before treatment. Cells were pre-treated with 1 μM arsenic for 24 hours before exposure to 3 kJ/m^2^ solar simulated ultraviolet light (ssUVR). Arsenic exposure was continued for 24 hours post ssUVR exposure and clones were expanded for DNA extraction and sequencing. DNA from each clone was extracted using the QIAamp® DNA Mini Kit (Qiagen).

### Cytotoxicity assay

Cytotoxicity was determined using a clonogenic survival assay modified from previously described methods^46^. Briefly, N/TERT1 cells were seeded and allowed to grow for 48 hours before treatment with 0 or 1 μM arsenic. After 24 hours, cells were exposed to increasing UVR doses. Cells were harvested immediately after UVR exposure and re-seeded in 100 mm dishes at colony forming density (300 cells/dish). After colony formation cells were fixed, stained with crystal violet, and colonies were counted. Four dishes per treatment group were included and results are expressed as relative survival, which was derived from the number of colonies per treatment group divided by the number of colonies in the control multiplied by 100.

### In vivo exposures and tissue collection

SKH-1 mice (21–25 days old) were purchased from Charles River Laboratories (Wilmington, MA). These studies were performed under an approved Institutional Animal Care and Use Committee (IACUC) protocol (#22-201244-HSC). Animals were housed by treatment group and administered arsenite (5 mg/l) in the drinking water for the duration of the study. Water was freshly prepared and changed every second day, and consumption monitored. There was equivalent water consumption between control and arsenic treated groups, and all animals were provided standard mouse chow ad lib. After 28 days of arsenic treatment, animals were exposed to UVR (14 kJ/m^2^; ∼0.5 minimal erythema dose [MED]) 3 times per week until the development of tumors (30 weeks). There were unavoidable UVR lamp issues during weeks 8 and 9 where animals were not UVR exposed. Water treatment continued for an additional 4 weeks to allow for tumor growth prior to collection. Tumor number by animal was determined once per week by physical palpation and counted if at least 1 mm in diameter. Some tumors regressed over time and only tumors that persisted for at least 3 weeks were included in the total count. Animals were euthanized using CO_2_ followed by cervical dislocation and tissues collected. Tissues collected included kidney, liver, spleen, ventral skin (UVR naïve), dorsal skin (UVR exposed) and skin tumors. Tissues were collected in 10% neutral buffered formalin, RNAlater, snap-frozen, and epidermal scrapings obtained from both ventral and dorsal skin

### DNA extraction from skin tumors and UVR naïve skin

Snap-frozen skin tumors (1.5 – 2 mm in diameter) and UV naïve skin sections (0.5 - 1 cm^2^) were thawed on ice then removed from the vial and placed on a glass plate previously cleaned with 70% ethanol and RNAzap (Thermo Fisher Scientific). Tissue was minced into small pieces with a pair of clean scalpels and transferred to clean RNase/DNase free tubes. Clean blades were used, and the mincing surface sanitized between samples to limit cross-contamination. Genomic DNA was extracted using the DNeasy Blood and Tissue kit (Qiagen) following the manufacturers recommendations. For the initial digestion step, 180 μl ATL buffer and 20 μl Proteinase K was added to each sample, vortexed thoroughly, then incubated at 50 °C for 2 hours with vortexing every 15 min. The remaining steps followed the kit’s directions exactly. DNA was eluted from the column with 2 consecutive additions of 50 μl of the AE buffer supplied with the kit. DNA concentration and quality was determined using Qubit (Thermo Fisher Scientific). Samples were subsequently diluted to the required concentrations for whole-genome sequencing.

### Whole-genome sequencing

DNA from *in vitro* and *in vivo* experiments was sent to Novogene (Sacramento, CA) and library preparation was performed using the NEBNext® DNA Library Prep Kit (New England Biolabs) following the manufacturer’s recommendations. Qualified libraries were sequenced on an Illumina platform to 30x coverage according to effective concentration and data volume.

### Identification of somatic mutations from whole genome bulk sequencing

Raw sequence data were downloaded to the Triton Shared Compute Cluster (TSCC) from ftp server link shared by Novogene (Sacramento, CA). All the post-sequencing analysis was performed within TSCC at UC San Diego. A schematics of the somatic mutations calling process is described in **Supplementary Fig. 5**. This methodology for identification of somatic mutations from bulk sequencing data follows established approaches from large genomics consortia^43^. Briefly, quality assurance of the raw FASTQ files were evaluated using FastQC and Mosdepth^47,48^. Raw sequence reads were aligned to the human reference genome GRCh38 for N/TERT1 data and GRCm39 for mouse data. The aligned reads were marked duplicated using MarkDuplicates (Picard) from GATK^49^. For human cell lines, concordance between exposed and stock samples were evaluated using Conpair^50^ and only samples with >99.5% concordance were taken forward for subsequent analysis. An ensemble variant calling pipeline (EVC) was used to identify single nucleotide variants (SNV) and short insertions and deletions (indels). EVC implements the SNV and indel variant calling from four variant callers (Mutect2, Strelka2, Varscan2, and MuSE) and only mutations that are identified by any two variant callers were considered as *bona fide* mutations^49,51-53^. For N/TERT1 cells, bulk sequencing data from stock were used as a matched normal. For, mouse data, ventral skin from each mouse was used as a matched normal.

### Analysis of mutational signatures

Analysis of mutational signatures was performed using our previously derived set of reference COSMIC mutational signatures^33^ as well as our previously established methodology with the SigProfiler suite of tools used for summarization, simulation, visualization, and assignment of mutational signatures. Briefly, mutational matrixes for SBS, DBS and ID were generated with SigProfilerMatrixGenerator^44^. Plotting of each mutational profile was done with SigProfilerPlotting. *De novo* mutational signature extraction and COSMIC decomposition of *de novo* signatures were performed with SigProfilerExtractor^54^. Attribution of COSMIC signatures to each of the samples mutational profile were performed using SigProfilerAssignment.

### Arsenic co-exposure validation in human cancer

To evaluate the potential arsenic exposure in human skin cancer through signature ID13, publicly available whole-genome sequenced skin melanomas and whole-exome sequenced basal cell carcinomas (BCCs) were evaluated. The mutational profiles and mutational signatures in each whole-genome sequenced melanoma were downloaded from a prior publication^22^. Whole-genome sequenced melanomas with at least 100 mutations contributing to ID13 were grouped as ID13 positive, while all remaining samples were classified as being ID13 negative. For whole-exome sequenced BCCs, somatic mutations were also derived from a prior publication^42^. Mutational signature extractions were performed using SigProfilerExtractor and samples containing ID13 were classified ID13 positive, while all remaining samples were classified as being ID13 negative.

Normalized mutational profiles and statistical significance testing were preformed within R statistical language^55^. Arrangements of figures and modifications were performed with Adobe Illustrator and BioRender^56^.

### Statistical analysis and reproducibility

All bar graphs are expressed as the mean ± SEM (standard error of the mean) with individual biological replicates shown as corresponding black circles. Since there are multiple distinct groups in the N/TERT1 experiments, one-way ANOVA with Tukey post hoc analysis for multiple comparisons was used to determine significance amongst controls and the samples in the three treatment groups. All cell culture groups have an *n*=3 except for the UVR alone group (*n*=2). Multiplicity adjusted p-values are reported with significance set to p-value<0.05 for all N/TERT1 analyses.

In the mouse study, whole-genome sequencing data were generated for the UVR group using 4 individual animals and for the arsenic plus UVR group using 3 individual animals. FDR-corrected two-sided t-tests were used to determine significance between UVR and arsenic plus UVR groups in all mouse analyses. FDR-corrections were performed using the Benjamini-Hochberg correction procedure. Significance was determined to be q-value<0.05 for all mouse analyses.

In human cancers, q-values were calculated using FDR-corrected two-sided pairwise t-tests. FDR-corrections were performed using the Benjamini-Hochberg correction procedure. Statistical significance was set at q-value<0.05. Statistical analysis and plotting were performed using GraphPad Prism v9.3.1.

### Code availability

Somatic mutations in whole-genome sequencing data were identified using our ensemble variant calling pipeline, which is freely available under the permissive 2-clause BSD license at: https://github.com/AlexandrovLab/EnsembleVariantCallingPipeline. All other computational tools utilized in this publication have been mentioned in the methodology section and can be access through their respective publications.

## Data availability

All whole-genome sequencing data have been deposited to Sequence Read Archive (SRA). The sequencing data for N/TERT1 cells can be downloaded using accession number: PRJNA909329 and for SKH-1 mice data with accession number: PRJNA91094. All data and metadata for the previously generated whole-genome sequenced melanoma cancers were obtained from the official PCAWG release (https://dcc.icgc.org/releases/PCAWG). Where appropriate, source data are provided for the figures in the paper

## Acknowledgements

Research reported in this publication was supported by the National Institute of Environmental Health Sciences [R01ES030993 to KJL/LGH], pilot grant (to LGH/KJL) and postdoctoral matching support (to RMS) by University of New Mexico Comprehensive Cancer Center through NIH/NCI grant [P30CA118100], and UNM Center for Metals in Biology and Medicine through NIH/NIGMS grant P20GM130422. Research performed for this publication at the Alexandrov Lab was supported by a Packard Fellowship for Science and Engineering as well as by grants from the US National Institutes of Health, including: NIEHS R01ES030993 (KJL/LGH), NIEHS R01ES032547 (LBA), and NCI R01CA269919 (LBA). The content is solely the responsibility of the authors and does not necessarily represent the official views of the National Institutes of Health. The funders were not involved in the study design, data collection, analysis and interpretation of the data, the writing of the article, or the decision to submit the article for publication.

## Author contributions

**RMS:** Conceptualization, Data curation, Formal analysis, Funding acquisition, Investigation, Methodology, Supervision, Validation, Visualization, Writing – original draft, Writing – review & editing. **SPN:** Conceptualization, Data curation, Formal analysis, Investigation, Methodology, Validation, Visualization, Writing – original draft, Writing – review & editing. **KLC:** Conceptualization, Data curation, Formal analysis, Investigation, Methodology, Validation, Visualization, Writing – original draft, Writing – review & editing. **XZ:** Conceptualization, Data curation, Formal analysis, Investigation, Methodology, Validation, Visualization, Writing – original draft, Writing – review & editing. **YG:** Conceptualization, Data curation, Formal analysis, Investigation, Methodology, Validation, Visualization, Writing – original draft, Writing – review & editing. **HY:** Conceptualization, Data curation, Formal analysis, Investigation, Methodology, Validation, Visualization, Writing – original draft, Writing – review & editing. **LGH:** Conceptualization, Data curation, Formal analysis, Funding acquisition, Investigation, Methodology, Project administration, Resources, Software, Supervision, Validation, Visualization, Writing – review & editing. **LBA:** Conceptualization, Data curation, Formal analysis, Funding acquisition, Investigation, Methodology, Project administration, Resources, Software, Supervision, Validation, Visualization, Writing – review & editing. **KJL:** Conceptualization, Data curation, Formal analysis, Funding acquisition, Investigation, Methodology, Project administration, Resources, Software, Supervision, Validation, Visualization, Writing – review & editing.

## REFERENCES

1 Barnes, J. L., Zubair, M., John, K., Poirier, M. C. & Martin, F. L. Carcinogens and DNA damage. Biochemical Society Transactions 46, 1213–1224 (2018).

2 Ames, B. N., Durston, W. E., Yamasaki, E. & Lee, F. D. Carcinogens are mutagens: a simple test system combining liver homogenates for activation and bacteria for detection. Proceedings of the National Academy of Sciences 70, 2281–2285 (1973).

3 Riva, L. et al. The mutational signature profile of known and suspected human carcinogens in mice. Nature genetics 52, 1189–1197 (2020).

4 Haverkos, H. W., Haverkos, G. P. & O’Mara, M. Co-carcinogenesis: Human papillomaviruses, coal tar derivatives, and squamous cell cervical cancer. Frontiers in microbiology 8, 2253 (2017).

5 Rossman, T. G., Uddin, A. N., Burns, F. J. & Bosland, M. C. Arsenite cocarcinogenesis: an animal model derived from genetic toxicology studies. Environmental Health Perspectives 110, 749–752 (2002).

6 Kitchin, K. T. Recent advances in arsenic carcinogenesis: modes of action, animal model systems, and methylated arsenic metabolites. Toxicology and applied pharmacology 172, 249–261 (2001).

7 Hughes, M. F., Kenyon, E. M. & Kitchin, K. T. Research approaches to address uncertainties in the risk assessment of arsenic in drinking water. Toxicology and applied Pharmacology 222, 399–404 (2007).

8 Hong, Y.-S., Song, K.-H. & Chung, J.-Y. Health effects of chronic arsenic exposure. Journal of preventive medicine and public health 47, 245 (2014).

9 Martinez, V. D., Vucic, E. A., Becker-Santos, D. D., Gil, L. & Lam, W. L. Arsenic exposure and the induction of human cancers. Journal of toxicology 2011 (2011).

10 Mostafa, M. G. & Cherry, N. Arsenic in drinking water, transition cell cancer and chronic cystitis in rural Bangladesh. International journal of environmental research and public health 12, 13739–13749 (2015).

11 Hartwig, A. et al. Mode of action-based risk assessment of genotoxic carcinogens. Archives of toxicology 94, 1787–1877 (2020).

12 Zhou, X., Speer, R. M., Volk, L., Hudson, L. G. & Liu, K. J. in Seminars in cancer biology. 86–98 (Elsevier).

13 Chen, Y. et al. Modification of risk of arsenic-induced skin lesions by sunlight exposure, smoking, and occupational exposures in Bangladesh. Epidemiology, 459–467 (2006).

14 Burns, F. J., Uddin, A. N., Wu, F., Nádas, A. & Rossman, T. G. Arsenic-induced enhancement of ultraviolet radiation carcinogenesis in mouse skin: a dose-response study. Environmental health perspectives 112, 599–603 (2004).

15 Ding, W., Hudson, L. G., Sun, X., Feng, C. & Liu, K. J. As (III) inhibits ultraviolet radiation-induced cyclobutane pyrimidine dimer repair via generation of nitric oxide in human keratinocytes. Free Radical Biology and Medicine 45, 1065–1072 (2008).

16 Rossman, T. G., Uddin, A. N. & Burns, F. J. Evidence that arsenite acts as a cocarcinogen in skin cancer. Toxicology and applied pharmacology 198, 394–404 (2004).

17 Chatterjee, N. & Walker, G. C. Mechanisms of DNA damage, repair, and mutagenesis. Environmental and molecular mutagenesis (2017).

18 Muenyi, C. S., Ljungman, M. & States, J. C. Arsenic disruption of DNA damage responses—potential role in carcinogenesis and chemotherapy. Biomolecules 5, 2184–2193 (2015).

19 Tam, L. M., Price, N. E. & Wang, Y. Molecular mechanisms of arsenic-induced disruption of DNA repair. Chemical research in toxicology 33, 709–726 (2020).

20 Alexandrov, L. B. et al. Signatures of mutational processes in human cancer. Nature 500, 415–421 (2013).

21 Alexandrov, L. B., Nik-Zainal, S., Wedge, D. C., Campbell, P. J. & Stratton, M. R. Deciphering signatures of mutational processes operative in human cancer. Cell reports 3, 246–259 (2013).

22 Alexandrov, L. B. et al. The repertoire of mutational signatures in human cancer. Nature 578, 94–101 (2020).

23 Martinez, V. D. et al. Whole-genome sequencing analysis identifies a distinctive mutational spectrum in an arsenic-related lung tumor. Journal of thoracic oncology 8, 1451–1455 (2013).

24 Kucab, J. E. et al. A compendium of mutational signatures of environmental agents. Cell 177, 821–836 (2019).

25 Hayward, N. K. et al. Whole-genome landscapes of major melanoma subtypes. Nature 545, 175–180 (2017).

26 Martincorena, I. et al. High burden and pervasive positive selection of somatic mutations in normal human skin. Science 348, 880–886 (2015).

27 Nik-Zainal, S. et al. The genome as a record of environmental exposure. Mutagenesis 30, 763–770 (2015).

28 Rawson, R. V. et al. Unexpected UVR and non-UVR mutation burden in some acral and cutaneous melanomas. Laboratory investigation 97, 130–145 (2017).

29 Saini, N. et al. The impact of environmental and endogenous damage on somatic mutation load in human skin fibroblasts. PLoS genetics 12, e1006385 (2016).

30 Pfeifer, G. P. & Besaratinia, A. UV wavelength-dependent DNA damage and human non-melanoma and melanoma skin cancer. Photochemical & photobiological sciences 11, 90–97 (2012).

31 Moreno, N. C. et al. Whole-exome sequencing reveals the impact of UVA light mutagenesis in xeroderma pigmentosum variant human cells. Nucleic acids research 48, 1941–1953 (2020).

32 Jin, S.-G., Padron, F. & Pfeifer, G. P. UVA Radiation, DNA Damage, and Melanoma. ACS omega (2022).

33 Tate, J. G. et al. COSMIC: the catalogue of somatic mutations in cancer. Nucleic acids research 47, D941–D947 (2019).

34 Dickson, M. A. et al. Human keratinocytes that express hTERT and also bypass a p16INK4a-enforced mechanism that limits life span become immortal yet retain normal growth and differentiation characteristics. Molecular and cellular biology 20, 1436–1447 (2000).

35 Zhivagui, M., Korenjak, M. & Zavadil, J. Modelling mutation spectra of human carcinogens using experimental systems. Basic & clinical pharmacology & toxicology 121, 16–22 (2017).

36 Cooper, K., Yager, J. & Hudson, L. Melanocytes and keratinocytes have distinct and shared responses to ultraviolet radiation and arsenic. Toxicology letters 224, 407–415 (2014).

37 Thompson, B. C., Halliday, G. M. & Damian, D. L. Nicotinamide enhances repair of arsenic and ultraviolet radiation-induced DNA damage in HaCaT keratinocytes and ex vivo human skin. PLoS One 10, e0117491 (2015).

38 Cooper, K. L. et al. Contribution of NADPH oxidase to the retention of UVR-induced DNA damage by arsenic. Toxicology and Applied Pharmacology 434, 115799 (2022).

39 Benavides, F., Oberyszyn, T. M., VanBuskirk, A. M., Reeve, V. E. & Kusewitt, D. F. The hairless mouse in skin research. Journal of dermatological science 53, 10–18 (2009).

40 Koller, B. H. et al. Arsenic metabolism in mice carrying a BORCS7/AS3MT locus humanized by syntenic replacement. Environmental health perspectives 128, 087003 (2020).

41 Vahter, M. Methylation of inorganic arsenic in different mammalian species and population groups. Science progress 82, 69–88 (1999).

42 Bonilla, X. et al. Genomic analysis identifies new drivers and progression pathways in skin basal cell carcinoma. Nature genetics 48, 398–406 (2016).

43 Pan-cancer analysis of whole genomes. Nature 578, 82–93 (2020).

44 Bergstrom, E. N. et al. SigProfilerMatrixGenerator: a tool for visualizing and exploring patterns of small mutational events. BMC genomics 20, 1–12 (2019).

45 Smits, J. P. et al. Immortalized N/TERT keratinocytes as an alternative cell source in 3D human epidermal models. Scientific reports 7, 1–14 (2017).

46 Wise Sr, J. P., Wise, S. S. & Little, J. E. The cytotoxicity and genotoxicity of particulate and soluble hexavalent chromium in human lung cells. Mutation Research/Genetic Toxicology and Environmental Mutagenesis 517, 221–229 (2002).

47 Andrews, S. (Babraham Bioinformatics, Babraham Institute, Cambridge, United Kingdom, 2010).

48 Pedersen, B. S. & Quinlan, A. R. Mosdepth: quick coverage calculation for genomes and exomes. Bioinformatics (Oxford, England) 34, 867–868 (2018).

49 McKenna, A. et al. The Genome Analysis Toolkit: a MapReduce framework for analyzing next-generation DNA sequencing data. Genome research 20, 1297–1303 (2010).

50 Bergmann, E. A., Chen, B.-J., Arora, K., Vacic, V. & Zody, M. C. Conpair: concordance and contamination estimator for matched tumor–normal pairs. Bioinformatics (Oxford, England) 32, 3196–3198 (2016).

51 Fan, Y. et al. MuSE: accounting for tumor heterogeneity using a sample-specific error model improves sensitivity and specificity in mutation calling from sequencing data. Genome biology 17, 1–11 (2016).

52 Kim, S. et al. Strelka2: fast and accurate calling of germline and somatic variants. Nature methods 15, 591–594 (2018).

53 Koboldt, D. C. et al. VarScan 2: somatic mutation and copy number alteration discovery in cancer by exome sequencing. Genome research 22, 568–576 (2012).

54 Islam, S. A. et al. Uncovering novel mutational signatures by de novo extraction with SigProfilerExtractor. Cell Genomics, 100179 (2022).

55 Team, R. C. R: A language and environment for statistical computing. (2013).

56 Perkel, J. M. The software that powers scientific illustration. Nature 582, 137–139 (2020).

