## Supplementary Figures for "Arsenic is a potent co-mutagen of ultraviolet light"

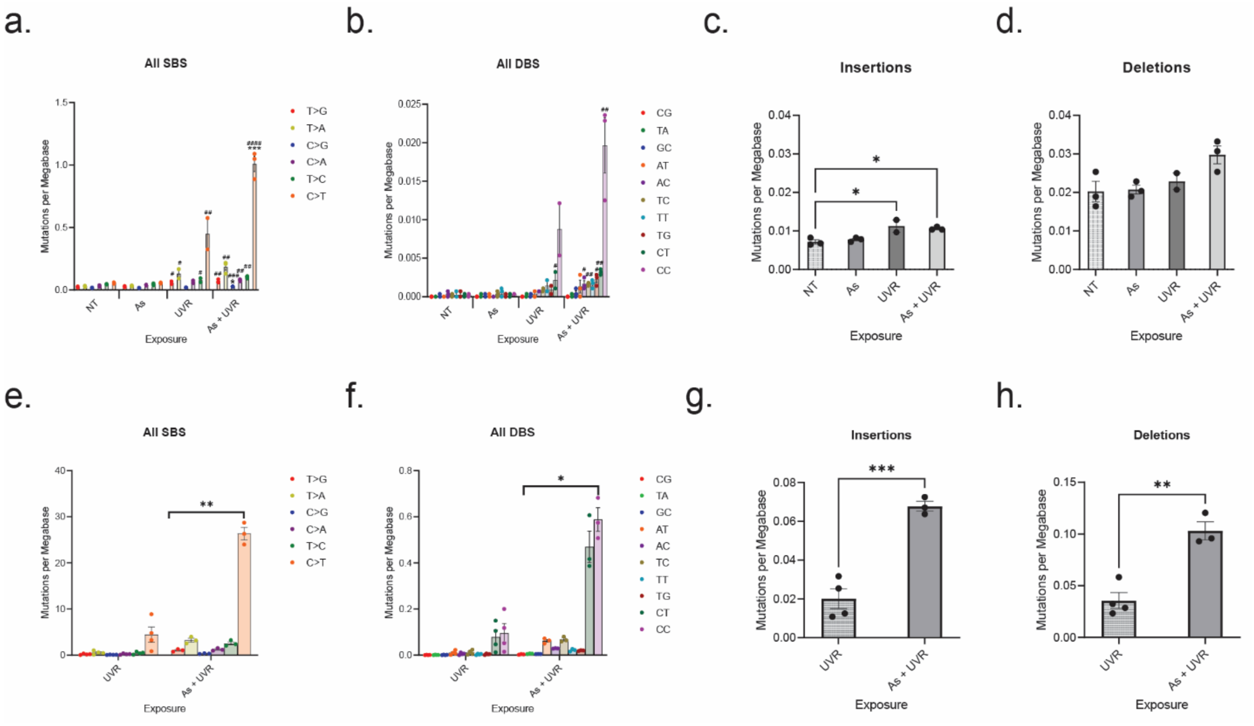


**Supplementary Figure 1. Arsenic enhances select UVR mutations. a.** In N/TERT1 cells UVR significantly increased T>G, T>A, C>A, T>C, and C>T mutations compared to the NT control and arsenic significantly enhanced C>G and C>T mutations compared to UVR alone. **b.** AC>NN, TC>NN, TG>NN, and CC>NN DBSs were significantly enhanced by arsenic and UVR co-exposure compared to the NT control in N/TERT1 cells. **c.** UVR and arsenic plus UVR co-exposure significantly increased insertions compared to the NT control, but not **d.** deletions. **e.** in SKH-1 tumors arsenic significantly increased all SBSs **f.** DBSs, **g.** insertions, and **h.** deletions. Significance was determined using one-way ANOVAs with Tukey's multiple comparisons test; n=3 for NT, As, and As plus UVR; n=2 for UVR. *p<0.05, **p-value<0.01, ***p-value <0.005. Bar plots represent the mean ± SEM; individual replicate values are shown as black circles. Statistical details are reported in the **Methods** section.

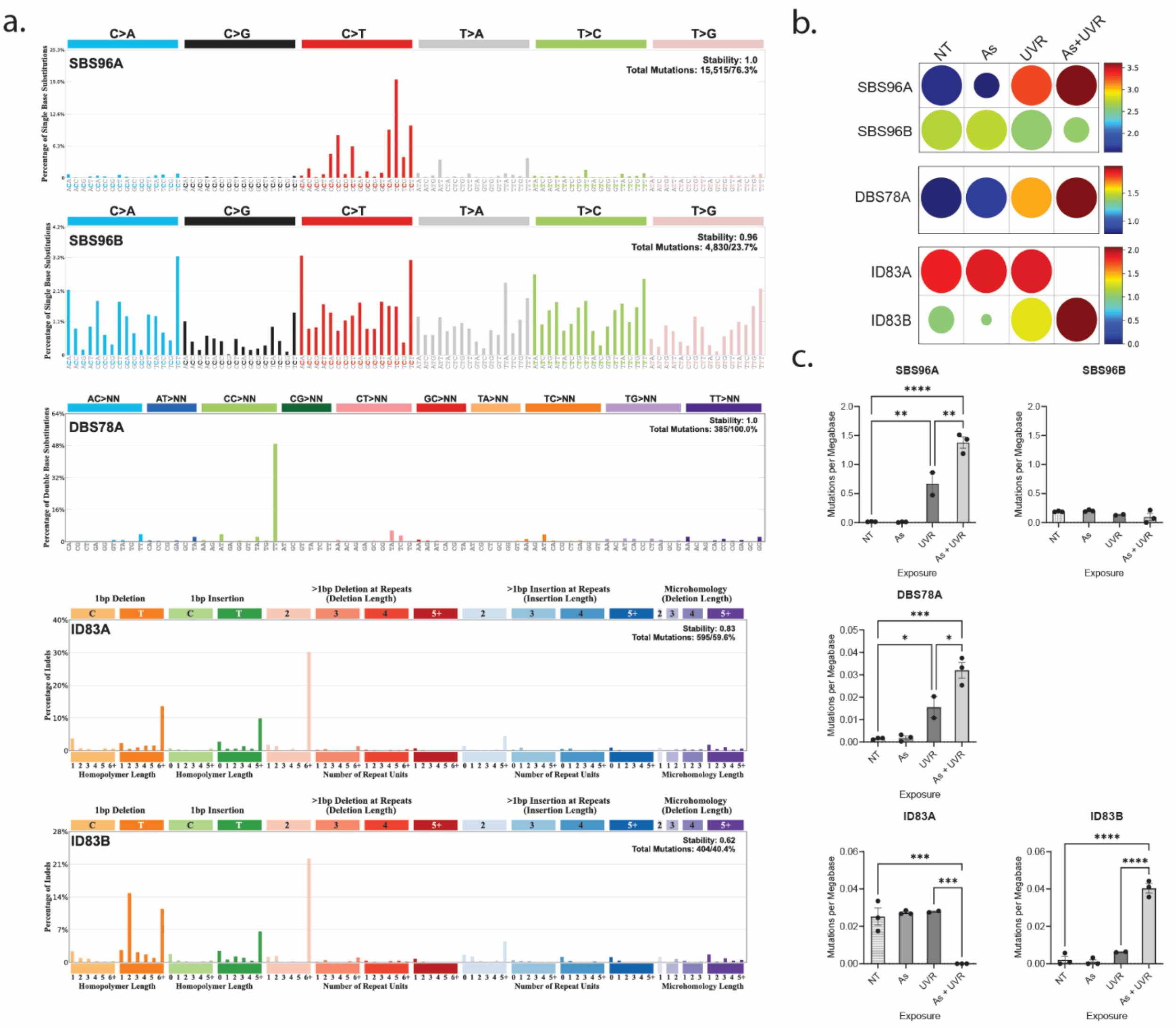


**Supplementary Figure 2. Analysis of *de novo* mutational signatures in N/TERT1 cells. a.** Mutational profiles of *de novo* signatures extracted in N/TERT1 cells including two SBS, one DBS, and two ID signatures. **b.** Contribution of *de novo* mutational signatures that underlie mutational profiles observed within N/TERT1 cells experiment. Each circle represents the activity of a signature for a given sample type. The radius of the circle determines the proportion of samples with greater than a given number of mutations specific to each subclass; the color reflects the $\log_{10}$ median number of mutations per treatment group. **c.** The mutations per megabase for each signature across treatment groups are shown. Significance was determined using one-way ANOVAs with Tukey's multiple comparisons test; n=3 for NT, As, and As plus UVR; n=2 for UVR. **p-value<0.01, ***p-value<0.005. Bar plots represent the mean ± SEM; individual replicate values are shown as black circles. Statistical details are reported in the **Methods** section.

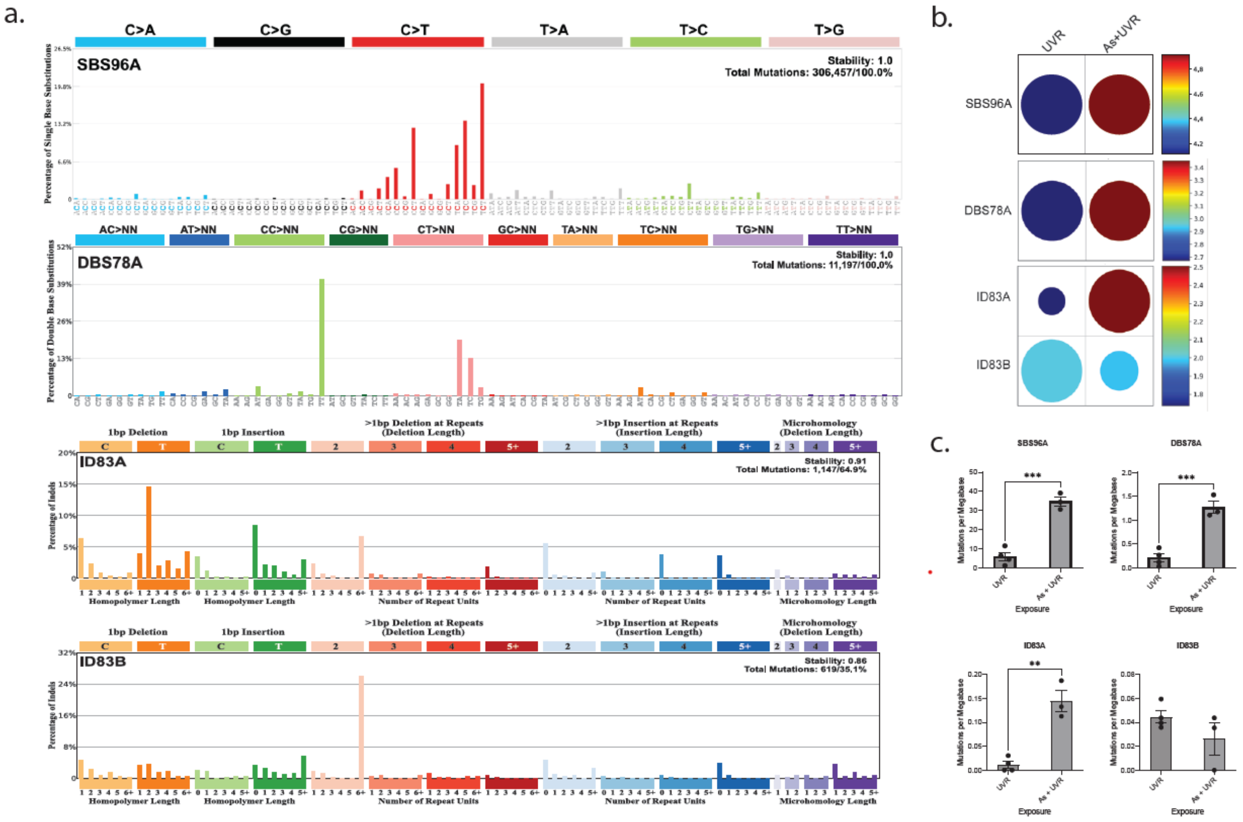


**Supplementary Figure 3.** ***De novo*** **mutational signatures in skin cancers from SKH-1 hairless mice. a.** Mutational profiles of *de novo* signatures extracted in SKH-1 tumors including one SBS, one DBS, and two ID signatures. **b.** Contribution of *de novo* mutational signatures that underlie mutational profiles observed within mouse tumor. Each circle represents the activity of a signature for a given sample type. The radius of the circle determines the proportion of samples with greater than a given number of mutations specific to each subclass; the color reflects the $\log_{10}$ median number of mutations per treatment group. **c.** The mutations per megabase for each signature across treatment groups are shown. Significance is determined using FDR-correcred unpaired 2-sided t-tests; n=4 for UVR alone and n=3 for As plus UVR derived. *q-value<0.05, ***q-value<0.05, ****q-value<0.001. Bar plots represent the mean ± SEM; individual replicate values are shown as black circles. Statistical details are reported in the **Methods** section.

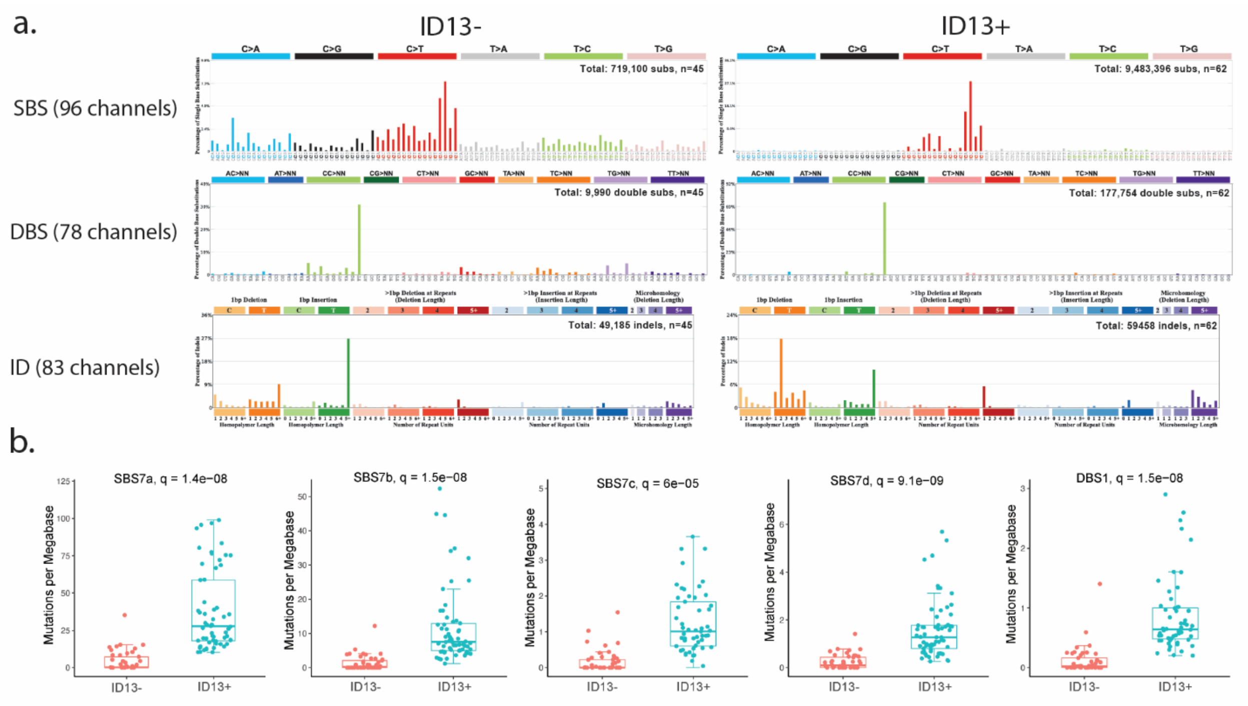


**Supplementary Figure 4. An evaluation of UVR and arsenic-like co-exposure in human skin melanomas. a.** Nominalized mutational profiles of single base substitutions, doublet base substitutions, and indels in their SBS-96, DBS-78 and ID-83 classificational schemas. Mutational profiles of ID13 negative and ID13 positive melanomas show similar SBS patterns characterized by C>T substitutions as well as similar DBS patterns characterized by CC>NN doublet substitutions. In contrast, a distinct difference is observed in the indel profiles designated by 1 base-pair thymidine deletions with a homopolymer length of 2, which are only observed in ID13 positive melanomas. **b.** Relative proportion to SBS7a, SBS7b, SBS7d, and DBS1 mutational signature per samples containing ID13 ($n$ = 62) or without ID13 ($n$ = 45; right and left, respectively) melanomas. Each dot reflects the mutations per megabase attributed to each COSMIC signature in each sample. The bounds of the boxplots represent the interquartile range divided by the median, and Tukey-style whiskers extend to a maximum of 1.5 × interquartile range beyond the box. Statistically significant results from FDR corrected two-sided t-tests tests are denoted as q-values.

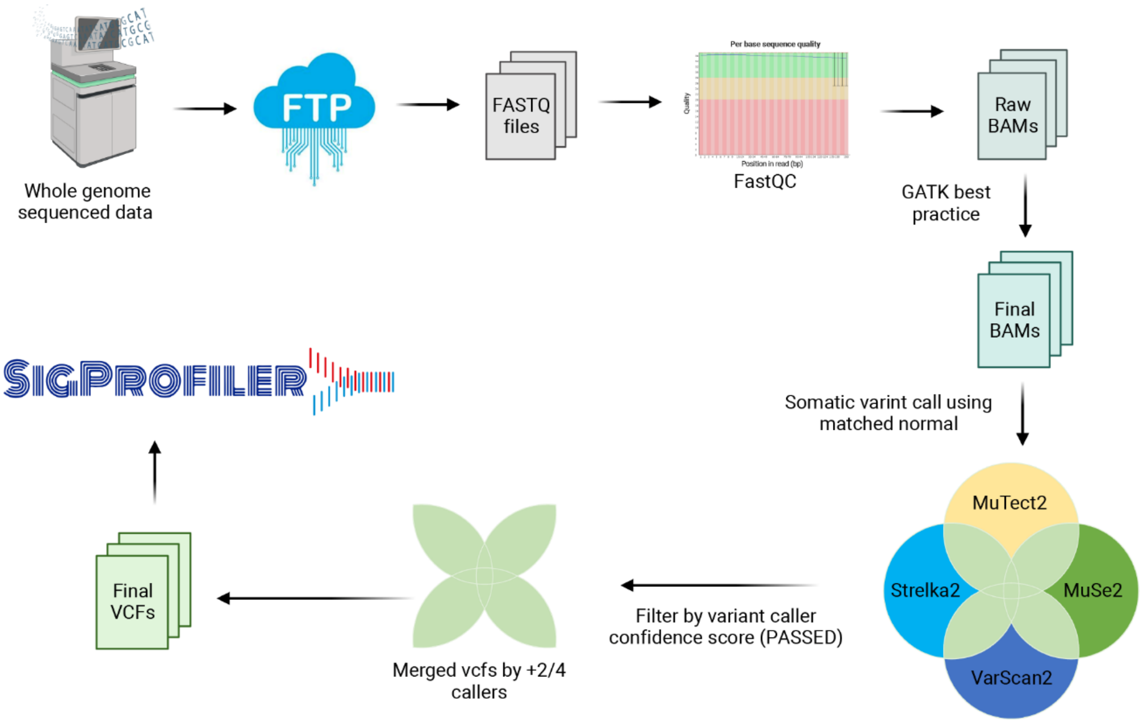


**Supplementary Figure 5. Schematic of whole-genome sequence data analysis.** Raw FASTQ files were downloaded within our shared computational cluster environment. Following GATK best practice, four variant callers (Mutect2, VarScan2, Strelka2, and MuSE) were employed in matched tumor-normal mode. Only mutations that are identified by any two variant callers were considered as *bona fide* mutations The final set of somatic mutations were analyzed by the SigProfiler suite of tools.
